# Intracellular dry mass density increases under growth-induced pressure

**DOI:** 10.1101/2024.09.10.612234

**Authors:** Hyojun Kim, Baptiste Alric, Nolan Chan, Julien Roul, Morgan Delarue

## Abstract

Cells that proliferate in confined environments develop mechanical compressive stress, referred to as growth-induced pressure, which inhibits growth and division across various organisms. Recent studies have shown that in these confined spaces, the diffusivity of intracellular nanoparticles decreases. However, the physical mechanisms behind this reduction remain unclear. In this study, we use quantitative phase imaging to measure the refractive index and dry mass density of *Saccharomyces cerevisiae* cells proliferating under confinement in a microfluidic bioreactor. Our results indicate that the observed decrease in diffusivity can be at least attributed to the intracellular accumulation of macromolecules. Furthermore, the linear scaling between cell content and growth-induced pressure suggests that the concentrations of macromolecules and osmolytes are maintained proportionally under such pressure in *S. cerevisiae*.

## 1. Introduction

Living cells proliferating in confined spaces eventually build up mechanical compressive stress exerted onto themselves and their surroundings. This growth-induced pressure (GIP) has the potential to decrease cell growth in all kingdoms of the living, including bacteria (1,2), fungi (3,4), plants (5,6) or mammals (7–9). It has recently been shown in the budding yeast *Saccharomyces cerevisiae* and in mammalian cells that GIP is accompanied by a decrease in the diffusion of genetically-encoded tracer nanoparticles (10). This decreased diffusion has been attributed to an increase in intracellular density through a mechanism where the production of macromolecules with limited cell volume expansion leads to increased biomass within the cytoplasm. However, the accumulation of macromolecules has yet to be observed experimentally, as this decrease in diffusion could also be attributed to other effects, such as a decrease in biochemical activity which is known to fluidize the cytoplasm (11,12).

The dry mass density of a living cell can be a direct proxy of macromolecular concentration. Numerous methods exist to measure the dry mass density of living cells, but their applications to confined cells developing GIP are inexistent. To develop GIP, cells are proliferating inside a confining microfluidic chamber and become densely packed (3,7). The experimental conditions restrict access to several measurement techniques, such as suspended microchannel resonator (SMR) measuring single cells’ buoyancy suspended in media (13,14), or cryoelectron tomography requiring cell fixation (15,16). On the other hand, quantitative phase imaging (QPI) is an alternative technique that can non-invasively measure the dry mass density of optically transparent biological cells or tissues, which does not require sample preparation (17,18).

QPI can quantify the refractive index (RI) distribution of a biological sample by detecting the phase difference of light passing through the sample with the surrounding media. This measured RI is directly proportional to the dry mass density of a biological sample (19). Here, we investigated the changes in the dry mass density of the budding yeast *Saccharomyces cerevisiae* growing within a confined space using QPI. We observed that confined growth was associated with increased dry mass density. We measured the concentration of a fluorophore inside the cell and observed that the increase in RI was proportional to the increase in fluorophore concentration. A linear extrapolation between GIP and RI down to the refractive index of water predicted the nominal intracellular osmotic pressure of the cell.

## 2. Methods and materials

### Cell culture conditions

*Saccharomyces cerevisiae* strains were grown and maintained on a Synthetic Complete (Formedium) + 2% dextrose (SCD) media agar Petri dishes. Next, a single colony was inoculated in a fresh liquid SCD liquid media and incubated with orbital shaking at 200 rpm at 30°C overnight. Exponentially growing culture at OD = 0.3 was then loaded into the microfluidic chamber.

### Microfabrication of PDMS devices

The mold with two layers of different heights is fabricated in a cleanroom at LAAS-CNRS using classical photolithography with negative photoresists. The first layer at a height of 0.8 μm defining the culture media channels was prepared with SU8 3000.5 (Kayaku Advanced Materials), and the second layer at a height of 9.2 μm defining the cell growth chamber and the main cell-loading channel was prepared with HARE-SQ10 (Kemlab). After the lithography process, perfluorodecyltrichlorosilane was grafted onto the wafer surface using an SPD (Memsstar Technology) machine to improve the hydrophobicity and non-stiction. PDMS elastomer was prepared by mixing the base and curing agent in a 10:1 ratio, poured onto the mold, and cured overnight at 60°C. Both PDMS and a glass coverslip were activated with oxygen plasma treatment (Diener PICO; gas, oxygen; pressure, 0.3 mbar; power, 100%; activation, 20 s), and bonded to each other immediately after surface activation. The PDMS/glass chip was then baked for ≥5 h at 60°C.

### Microfluidic device operation

The cell suspension was introduced to the chambers using a syringe. We next injected the SCD media through the main inlet channel using a Fluigent MFCS pressure control system. During the cell incubation and imaging, the pressure at the culture media inlet was maintained at ∼1 bar. The chips were placed in a microscope environmental chamber at 30°C (Leica TempControl 37).

### Refractive index measurement of bulk liquid/solid samples

Liquid and solid samples were measured using an Abbe refractometer (OPL, France) with white light at 30°C. Culture media (SCD) and distilled water samples were measured in spread, like a thin film, between the main and secondary prisms. A cured PDMS sample was prepared into a film of about 500 μm thickness and was carefully placed between the two prisms’ surfaces without any air bubbles. The RI of each sample was consistent for three independent measurements, and errors were not stated because the variances were smaller than the refractometer’s measurement resolution (10^−4^).

### Bright field and fluorescent microscopy

All imaging acquisition in this study was conducted on a Leica DMi8 microscope. The lateral deformation of PDMS chambers was measured to infer growth-induced pressure through bright-field microscopy simultaneously with the QPI. The relationship between pressure and chamber deformation was calibrated as done in previous work (10), giving a value of 8.2 μm.MPa^−1^.

To measure GFP accumulation, cytosolic GFP expressed from *HIS3* promoter were measured by fluorescent z-stacking (0.5 μm interval) with a spinning-disk confocal scanner unit (Yokogawa CSU-X1). We acquired z-stack of the first cell layer of each chamber using the same way of measurement for the chamber height with ET525/50 nm red emission filter, a dichroic mirror (ZT405/488/561/638rpc, Chroma) and Hamamatsu scientific complementary metal–oxide– semiconductor camera (ORCA-Flash4.0 v3). In the same way, we imaged the cells without any GFP-labeling to subtract the cellular autofluorescence which can be detected from GFP channels. The GFP expression of each cell was calculated by summing of pixel values across all z-stack slice of an chamber in ImageJ/Fiji (20).

### Refractive index and dry mass density measurement of cells with QPI

For transmission QPI, the microscope was equipped with conventional LED Köhler transillumination and a condenser of maximum numerical aperture NA = 0.55. To acquire QPI data, we imaged the samples with a 20x 0.8 NA objective (Leica) and a quadriwave lateral shearing interferometry (QWLSI) system (SID4 sc8, Phasics) mounted on the microscope’s lateral camera port. The QWLSI measures the local phase shift, also called optical path difference (OPD), introduced by a specimen placed under a microscope. Depending on the sample’s thickness and the difference of sample’s refractive index from the background material, the OPD was depicted in grayscale on the image, the so-called phase image.

To estimate RI of cells (n_cell_) on a culture plate, every cell was segmented from a phase image, which set the culture media as a background baseline, using Otsu algorithm method by CellProfiler (20). The n_cell_ is estimated by the relation, n_cell_ = n_background_ + OPD/d_cell_, where the OPD is difference between cell and background and d_cell_ is the height of the cell. Since cells were assumed to be prolate ellipsoids, the d_cell_ was taken as the length of the minor axis of the cell in the phase image. The corresponding OPD was taken as the brightness of the upper 5% of the intensity distribution within the cell from the phase image in which the background was subtracted.

To measure the average RI of cells within a microfluidic chamber, the averaged OPD value along the chamber area was measured, comparing to surrounding PDMS. To minimize the influence of local and variable OPD gradients occurring in phase images of cells within the PDMS microfluidics chip, we took the average value of the PDMS area in all directions surrounding the cell sample as the background baseline of the measurement. The chamber height under pressure was determined by the OPD of the calibrated culture medium of chambers expanded by predefined hydraulic pressure: d_chamber_= OPD/(n_PDMS_-n_medium_). Measurement of chamber’s height at GIP = 0 MPa perfectly matched the measurement of the SU8 mold measured with a profilometer. For GIP > 0 MPa, the cellular RI was measured in comparison to PDMS: n_cell_(P) = n_PDMS_ + OPD(P)/d_chamber_(P).

The dry mass density of a yeast cell is directly calculated from the mean RI value since the RI in biological samples is linearly proportional to the dry mass density inside cells as n_cell_ = n_media_ + αρ, where n_cell_ is the average RI of each cell, n_media_ is the RI of the surrounding media which was obtained with Abbe refractometer (n_media_ = 1.336), α is an RI increment (α = 0.190 ml/g for proteins and nucleic acids(19)), and ρ is the dry mass density inside cells.

### Data and statistical analysis

All image analysis was performed using ImageJ (National Institutes of Health, US). Statistical analysis was performed on Python with Statsmodels modules (21).

## 3. Results

### Refractive index and dry mass density of S. cerevisiae determined with QPI

We measured the RI and dry mass density of living yeast *S. cerevisiae* on a culture plate using QPI, which measures the phase difference between a sample and a reference in the image. Phase images displayed the optical path difference (OPD) between cells and the surrounding culture media in grayscale (Fig. 1a). The OPD was the difference in RI of a sample with regard to a reference (here, the culture media), times the height of the sample. The pixel value along the cross section of a cell was related to the physical thickness and the local RI of different parts of living yeasts (Fig. 1b). We measured the maximum OPD of each cell with respect to the culture media to be 180 ± 20 nm. We estimated the average cellular height to be 3.94 ± 0.50 μm (see Materials and Methods). We determined the mean RI and dry mass density of an asynchronous population of yeast *S. cerevisiae*, n_cell_ = 1.384 ± 0.004 and ρ_cell_ = 271 ± 19 mg/ml (Fig. 1c), which lied within the range of the RI measured for various yeast cells (22–26).

**Figure 1.**
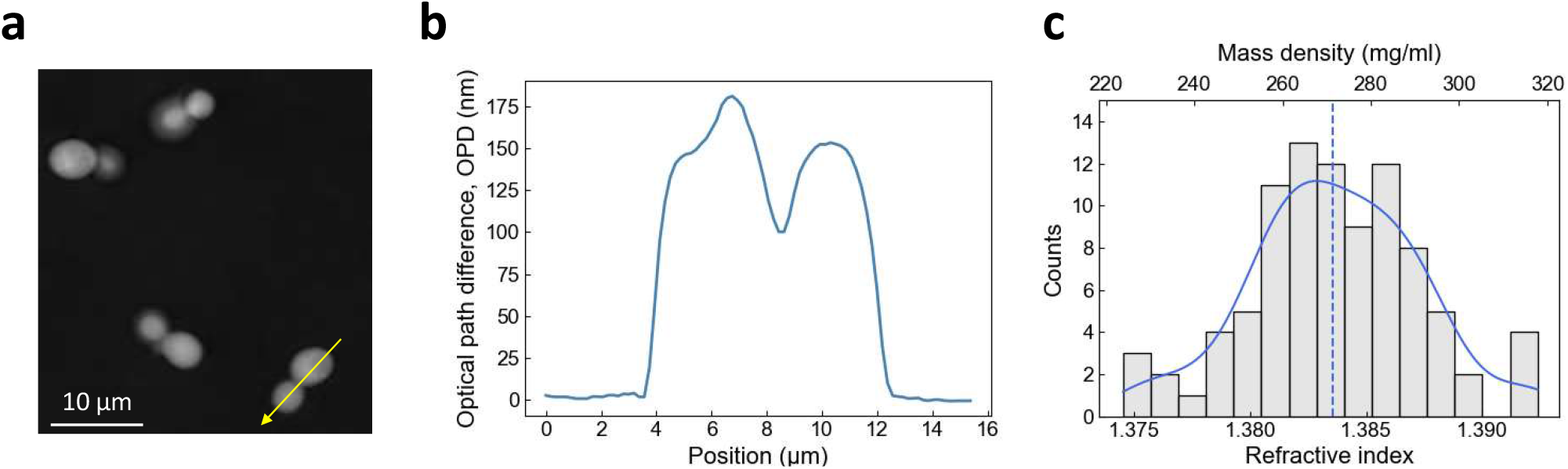
Measurement of cellular refractive index and dry mass density using QPI. (a) Optical path difference (OPD) image of budding yeast cells in SCD medium. (b)Profile plot drawn from (a) following the yellow arrow line. (c)Histogram of Refractive index (RI) and dry mass density measurements of budding yeast cells grown at 30°C in SCD medium. A kernel density estimation was plotted (blue) with the median value (dashed line).

### Increased intracellular density under GIP

We investigated the change in intracellular dry mass density as a function of growth-induced pressure (GIP). We used microfluidic elastic chambers to grow *S. cerevisiae* within a confined space to investigate the cellular RI under GIP (Fig. 2a). To prevent nutrient depletion and exchange media, chambers were supplied with nutrients through microchannels on both sides. After the cells filled the space through proliferation, they developed GIP by pushing against their neighbors and onto their surroundings. GIP was measured through the deformation of the PDMS elastic wall.

**Figure 2.**
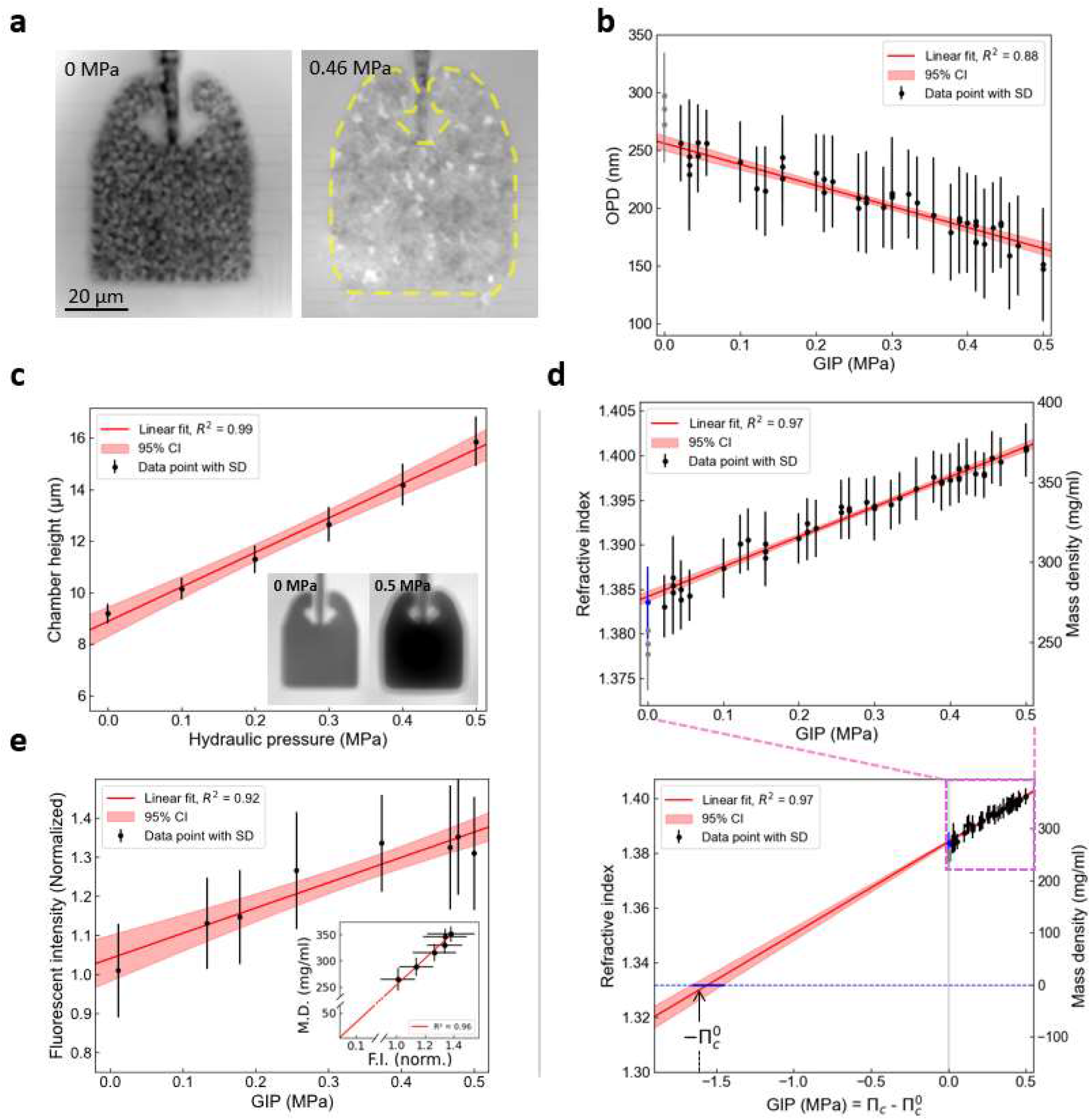
Intracellular dry mass density of budding yeast increases linearly with growth-induced pressure. (a)OPD images of budding yeast cells growing within the confining space of a PDMS microfluidic chamber. Cells do not experience growth-inducing pressure until their growth fills the entire volume within the chamber (left, GIP = 0 MPa). After, confined growth leads to the build-up of growth-induced pressure (GIP), deforming the elastic chamber, where the expanded boundary is labeled with a yellow dashed line (right, GIP = 0.46 MPa). (b) Change in OPD of cells under GIP measured relative to surrounding PDMS. (c) The effective height of the PDMS chamber linearly increased with the GIP level, which is proportional to the OPD between the chamber and surrounding PDMS (inset). (d) The RI and dry mass density increase linearly as a function of GIP. The value estimated outside of the device previously is presented in a blue point. The measurements within the device at GIP = 0 are presented in grey, which were excluded from the regression as they are assumed to be underestimated. The nominal intracellular osmotic pressure 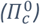 is estimated through linear extrapolation. An estimated 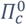value is indicated. (e) Fluorescence intensity (F.I.), corresponding to GFP expression level, linearly increases along GIP (n = 20). Dry mass (D.M.) density is proportional to fluorescence intensity (inset). The values in the inset graph is denoted by mean ± standard deviation in both the x and y-axis direction.

We acquired phase images of cells in a chamber under GIP using QPI to examine cellular RI (Fig. 2a). We measured that the average OPD of the cells with respect to the surrounding PDMS decreased linearly with pressure (Fig. 2b). As PDMS has a higher RI than the cells on average, n_PDMS_ = 1.405, the decrease in OPD implied that the RI of a cell approached the value of PDMS with increased GIP. We measured the effective height of the deformed confining chamber using OPD of the chamber filled with calibrated culture medium and pressurized by a measured hydraulic pressure (Fig. 2c). We then extracted, assuming that the chamber was fully filled with cells, the mean cellular RI as a function of GIP (Fig. 2d). We showed that RI increased roughly linearly with increased GIP under confinement.

We noted that the RI of cells without pressure in the chambers was underestimated (gray points in Fig. 1c, compared to the blue point measured outside of the device). We attributed this underestimation to the fact that cells were not tightly packed in the chamber, lowering the effective RI due to culture medium at a non-negligeable volume fraction in the chamber.

### Proportional increase in GFP production with intracellular density under GIP

We measured the mean fluorescence intensity of a GFP expressed from the *HIS3* promoter. We showed a linear increase of GFP concentration in the cell as a function of GIP (Fig. 2e). Interestingly, we observed that the increase in mass density was proportional to the increase in GFP concentration (Fig. 2e, inset), suggesting that the linearity between RI and protein concentration was kept in these conditions. Importantly, intracellular density extrapolated to 0 when the fluorescence intensity did. These results together demonstrated that the cellular biomass and GFP concentration increase roughly proportionally to GIP.

### Estimation of the nominal intracellular osmotic pressure

The intracellular RI is proportional to the dry mass of the cell. Fig. 2d showed that mass and intracellular osmotic pressure increased proportionally under confined growth. We performed a linear extrapolation of the pressure to a RI matching the one of water at 30°C (n_water_ = 1.332), without considering the points at GIP = 0 MPa measured in the microfluidic chip, but considering the RI of cells measured in culture medium (blue point). We assumed that when the cell would have the RI of water, it would be empty of its constituents, and its intracellular pressure would then be null, *Π*_*c*_ = O MPa. When the RI would match the RI of water, it would thus correspond to a point where the cell would be “empty” of macromolecules. The corresponding GIP, denoted 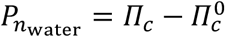, corresponding to the pressure difference between the intracellular osmotic pressure cell, *Π*_*c*_, and the nominal intracellular osmotic pressure, 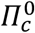, allowed us to estimate the nominal intracellular pressure of the cell: 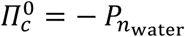 (Fig. 2d). We found in this case a nominal pressure of 1.55 MPa and the 95% confidence interval ranges from 1.46 to 1.66 MPa, comparably larger than the 0.95 MPa estimated for the same *S. cerevisiae* in previous studies, with a different method (10) (see Discussion).

## 4. Conclusion

In this study, the dry mass density under pressure was estimated by quantitative phase microscopy, and increased with GIP. The previous measurement of intracellular density under GIP has only been estimated indirectly by tracking fluorescently tagged tracer nanoparticles. It was assumed that the decrease in particle diffusion occurred due to the reduction of the free volume of the cytoplasm (10). However, the macromolecular diffusion within the cytoplasm, a highly heterogeneous active media, can be limited not only by the accumulation of macromolecules but also by a change in macromolecule size distribution (27), molecular electrostatic interactions (28), or active random force (29). Therefore, our results experimentally verify that the physical basis of the crowding-induced reduction in particle diffusion is compatible with an increase in dry mass density.

The previous theoretical model relating nanoparticle tracer diffusion with GIP assumed that the concentrations of osmolytes and macromolecules were produced proportionally (10,33), but this was not confirmed experimentally. The linear relation between GIP and RI shown in our results indicated that the production ratio of macromolecules and osmolytes was kept constant under pressure in the yeast *S. cerevisiae* (Fig 2d). However, the nominal osmotic pressure of budding yeast estimated through linear extrapolation, 1.55 MPa, was slightly larger than the value obtained from the previous study’s theoretical model, 0.95 MPa (10), but remained in the same order of magnitude.

## 5. Data and Software Availability

All acquired data and the codes developed in this work are freely available.

## 6. Author Contributions

HK, BA, NC, JR and MD did the experiments. HK and MD analyzed the data and wrote the manuscript. All authors proofread the manuscript.

## 7 Acknowledgments

Technological realizations were partly supported by the French RENATECH network. This work is partly funded by the European Union (ERC, UnderPressure, grant agreement number 101039998). Views and opinions expressed are however those of the author(s) only and do not necessarily reflect those of the European Union or the European Research Council. Neither the European Union nor the granting authority can be held responsible for them.

## 8. Conflict of interests

None to declare.

## Notes

### Competing Interest Statement

The authors have declared no competing interest.

## Reference

(1) Stewart PS, Robertson CR. Microbial growth in a fixed volume: studies with entrapped Escherichia coil. Applied Microbiology and Biotechnology. 1989;30:34–40. doi: 10.1007/BF00255993

(2) Chu EK, Kilic O, Cho H, Groisman A, Levchenko A. Self-induced mechanical stress can trigger biofilm formation in uropathogenic Escherichia coli. Nat Commun. 2018;9(1):4087. doi:10.1038/s41467-018-06552-z

(3) Delarue M, Hartung J, Schreck C, et al. Self-driven jamming in growing microbial populations. Nature Phys. 2016;12(8):762–766. doi:10.1038/nphys3741

(4) Delarue M, Poterewicz G, Hoxha O, et al. SCWISh network is essential for survival under mechanical pressure. Proceedings of the National Academy of Sciences. 2017;114(51):13465–13470. doi:10.1073/pnas.1711204114

(5) Bengough AG, Croser C, Pritchard J. A biophysical analysis of root growth under mechanical stress. In: Anderson HM, Barlow PW, Clarkson DT, Jackson MB, Shewry PR, eds. Plant Roots -From Cells to Systems. Springer Netherlands; 1997:107–116. doi:10.1007/978-94-011-5696-7_11

(6) Breuil-Broyer S, Morel P, De Almeida-Engler J, Coustham V, Negrutiu I, Trehin C. High-resolution boundary analysis during Arabidopsis thaliana flower development. The Plant Journal. 2004;38(1):182–192. doi:10.1111/j.1365-313X.2004.02026.x

(7) Meriem ZB, Mateo T, Faccini J, et al. A microfluidic mechano-chemostat for tissues and organisms reveals that confined growth is accompanied with increased macromolecular crowding. Lab Chip. 2023;23(20):4445–4455. doi:10.1039/D3LC00313B

(8) Rizzuti IF, Mascheroni P, Arcucci S, et al. Mechanical Control of Cell Proliferation Increases Resistance to Chemotherapeutic Agents. Phys Rev Lett. 2020;125(12):128103. doi:10.1103/PhysRevLett.125.128103

(9) Alessandri K, Sarangi BR, Gurchenkov VV, et al. Cellular capsules as a tool for multicellular spheroid production and for investigating the mechanics of tumor progression in vitro. Proc Natl Acad Sci U S A. 2013;110(37):14843–14848. doi:10.1073/pnas.1309482110

(10) Alric B, Formosa-Dague C, Dague E, Holt LJ, Delarue M. Macromolecular crowding limits growth under pressure. Nat Phys. 2022;18(4):411–416. doi:10.1038/s41567-022-01506-1

(11) Weber SC, Spakowitz AJ, Theriot JA. Nonthermal ATP-dependent fluctuations contribute to the in vivo motion of chromosomal loci. Proceedings of the National Academy of Sciences. 2012;109(19):7338–7343. doi:10.1073/pnas.1119505109

(12) Sakaue T, Saito T. Active diffusion of model chromosomal loci driven by athermal noise. Soft Matter. 2016;13(1):81–87. doi:10.1039/C6SM00775A

(13) Grover WH, Bryan AK, Diez-Silva M, Suresh S, Higgins JM, Manalis SR. Measuring single-cell density. Proceedings of the National Academy of Sciences. 2011;108(27):10992–10996. doi:10.1073/pnas.1104651108

(14) Miettinen TP, Ly KS, Lam A, Manalis SR. Single-cell monitoring of dry mass and dry mass density reveals exocytosis of cellular dry contents in mitosis. Pines J, Cooper JA, Piel M, eds. eLife. 2022;11:e76664. doi:10.7554/eLife.76664

(15) Delarue M, Brittingham GP, Pfeffer S, et al. mTORC1 Controls Phase Separation and the Biophysical Properties of the Cytoplasm by Tuning Crowding. Cell. 2018;174(2):338-349.e20. doi:10.1016/j.cell.2018.05.042

(16) Asano S, Engel BD, Baumeister W. In Situ Cryo-Electron Tomography: A Post-Reductionist Approach to Structural Biology. Journal of Molecular Biology. 2016;428(2, Part A):332–343. doi:10.1016/j.jmb.2015.09.030

(17) Park Y, Depeursinge C, Popescu G. Quantitative phase imaging in biomedicine. Nature Photon. 2018;12(10):578–589. doi:10.1038/s41566-018-0253-x

(18) Nguyen TL, Pradeep S, Judson-Torres RL, Reed J, Teitell MA, Zangle TA. Quantitative Phase Imaging: Recent Advances and Expanding Potential in Biomedicine. ACS Nano. 2022;16(8):11516–11544. doi:10.1021/acsnano.1c11507

(19) Zhao H, Brown PH, Schuck P. On the Distribution of Protein Refractive Index Increments. Biophysical Journal. 2011;100(9):2309–2317. doi:10.1016/j.bpj.2011.03.004

(20) Stirling DR, Swain-Bowden MJ, Lucas AM, Carpenter AE, Cimini BA, Goodman A. CellProfiler 4: improvements in speed, utility and usability. BMC Bioinformatics. 2021;22(1):433. doi:10.1186/s12859-021-04344-9

(21) Seabold S, Perktold J. Statsmodels: Econometric and Statistical Modeling with Python. In: ; 2010:92–96. doi:10.25080/Majora-92bf1922-011

(22) Abuhattum S, Kim K, Franzmann TM, et al. Intracellular Mass Density Increase Is Accompanying but Not Sufficient for Stiffening and Growth Arrest of Yeast Cells. Frontiers in Physics. 2018;6. Accessed January 28, 2024. https://www.frontiersin.org/articles/10.3389/fphy.2018.00131

(23) Odermatt PD, Miettinen TP, Lemière J, et al. Variations of intracellular density during the cell cycle arise from tip-growth regulation in fission yeast. Balasubramanian MK, Barkai N, Piel M, eds. eLife. 2021;10:e64901. doi:10.7554/eLife.64901

(24) Bianco V, D’Agostino M, Pirone D, et al. Label-Free Intracellular Multi-Specificity in Yeast Cells by Phase-Contrast Tomographic Flow Cytometry. Small Methods. 2023;7(11):2300447. doi:10.1002/smtd.202300447

(25) Boltyanskiy R, Odete MA, Cheong FC, Philips LA. Label-free viability assay using in-line holographic video microscopy. Sci Rep. 2022;12(1):12746. doi:10.1038/s41598-022-17098-y

(26) Midtvedt D, Olsén E, Höök F, Jeffries GDM. Label-free spatio-temporal monitoring of cytosolic mass, osmolarity, and volume in living cells. Nat Commun. 2019;10(1):340. doi:10.1038/s41467-018-08207-5

(27) Kalwarczyk T, Tabaka M, Holyst R. Biologistics—Diffusion coefficients for complete proteome of Escherichia coli. Bioinformatics. 2012;28(22):2971–2978. doi:10.1093/bioinformatics/bts537

(28) Zhou HX, Pang X. Electrostatic Interactions in Protein Structure, Folding, Binding, and Condensation. Chem Rev. 2018;118(4):1691–1741. doi:10.1021/acs.chemrev.7b00305

(29) Brangwynne CP, Koenderink GH, MacKintosh FC, Weitz DA. Cytoplasmic diffusion: molecular motors mix it up. Journal of Cell Biology. 2008;183(4):583–587. doi:10.1083/jcb.200806149

